# A spoonful of what helps the medicine go down? Improving the reliability of voluntary ingestion for oral dosing in rats and mice

**DOI:** 10.64898/2026.03.04.709533

**Authors:** Julia Bartlett, Emma S J Robinson

## Abstract

Voluntary ingestion is a refined method for substance administration that can replace oral gavage in rats and mice. It requires no physical restraint and has no associated risks of adverse effects, resulting in improved welfare and reduced distress for both animals and research staff. This method has been shown to be effective for a variety of compounds but is still not widely used due to concerns about accuracy and reliability. One potential issue is aversion to the taste of the compound being administered, including a common issue of bitter taste. In this study we tested compounds used in oral preparations for human medicines to mask bitter tasting drugs, including a commercial formulation designed for this purpose, Bitter Drug Powder™ (BDP). The masking agents were given in combination with a palatable vehicle (10% condensed milk) and the amount consumed and time to consume recorded. Animals were first habituated to the vehicle with reliable ingestion achieved within a few days. In the first studies, only BDP was fully effective at masking the bitter taste of quinine and preventing the progressive reduction in reliability of intake of the antidepressant, venlafaxine in mice and rats. We were able to replicate these effects using a combination of two different artificial sweeteners, saccharine and acesulfame K, and a thickening agent xanthan gum. These studies demonstrate that using a masking agent can improve the reliability of voluntary oral dosing in mice and rats and provide evidence to support a formulation which is readily available for researchers.

## Introduction

Animal studies frequently involve the administration of test substances using oral gavage. This method requires physical restraint which causes distress to the animal (1,2). This negatively impacts welfare (3,4) and scientific outcomes (5,6,7,8). The stress caused to the experimenter is detrimental to the emotional welfare of staff and can increase the risk of compassion fatigue (9). In addition to causing stress, this method can be physically damaging to the animal as incorrect placement of the gavage catheter can lead to tracheal dosing and oesophageal trauma (11,12,13). An alternative method for oral dosing is voluntary ingestion. This method involves combining the test compound with a palatable vehicle. This removes the risks of oral gavage and physical restraint, eliminating stress to the animal and experimenter. Voluntary ingestion has been shown to produce equivalent plasma concentrations of the administered drug when compared to oral gavage (14,15,16). There have been many groups that have successfully used voluntary ingestion in their studies (17,18,19,20,21) but the uptake of these techniques has not been widespread despite its welfare benefits. It is not clear why this refinement has not become more widely adopted but we suspect one factor may be concerns about the reliability and accuracy of these methods or issues which have arisen when trying the method. Within our lab, we have regularly used this technique and generated a protocol that improves the reliability and accuracy (22). Our protocol involves first habituating the animals to drinking a palatable vehicle from a syringe to overcome the neophobia that rodents can display when encountering a novel food (23,24). It also allows for drugs to be formulated to exact concentrations and, as the dose is presented in a syringe or micropipette (25), an exact volume can be administered. Animals can be dosed quickly using this method and are observed throughout the procedure so we can be confident that the entire dose has been consumed. The syringe/pipette-feeding method results in animals that will reliably and quickly consume test compounds when presented (18,26). However, there are some test compounds that animals either refuse to consume or gradually consume less of over repeated presentations (18,27), which may be related to taste aversion, particularly as many drugs are bitter.

Taste aversion is a similar challenge for paediatric and geriatric medicine where patients unable to swallow pills require oral formulations. Work conducted in these fields has identified compounds which improve palatability of aversive tasting medicines (28,29,30). Our aim was to investigate whether any of these compounds would be effective for masking taste for rodent voluntary ingestion. The masking compounds selected initially were saline, saccharine, and sodium gluconate, all of which have been shown to mask bitter tastes (30,31). These were tested using the bitter substance quinine which is used in human studies of taste masking and reliably decreases voluntary consumption without any adverse effects to the human or animal (32,31). Through our research into taste-masking technologies, we also discovered a compound developed by PCCA ltd called Bitter Drug Compound™ (BDP). We tested the effectiveness of BDP on quinine consumption in mice and rats and the reliability of repeated venlafaxine administration in rats. Previous studies with venlafaxine had found that, while animals would consume the drug on the first dosing day, the reliability would reduce over repeated administrations. As BDP is a propriety formulation and most researchers require more specific information about the vehicle formulation, we also developed our own taste masking compound using two sweetening agents along with a thickening agent. This formulation was based on human studies suggesting this is an effective strategy for creating a palatable liquid formulation for oral ingestion (33). Studies in rats also suggest combining a sweet tasting artificial flavouring with another artificial sweetener was effective in masking an aversive taste (34). Following pilot testing of different combinations of various agents, we settled on a 1:1:1 mixture of saccharine, acesulfame K and xanthan gum and tested the efficacy of this masking mixture (MM) alongside BDP at masking the bitter taste of quinine for both acute and repeated administration.

## Methods

### Subjects

These studies used 5 Cohorts of male lister hooded rats, 2 cohorts of CD1 mice and 1 cohort of 127sv, C57bl and BALBC male mice (Table 1). All animals were housed in temperature-controlled conditions ∼21°C on a 12:12 light cycle (lights off at 08:15 am). All dosing was conducted during the animal’s active phase. The rats were housed in pairs in conventional, open-top NKP large rat cages with a red Perspex platform, cardboard tube, wooden block, aspen ball, and rope. The mice were either single housed or pair housed, in conventional, open-top Techniplast 1284 caging with a carboard house, wooden block, aspen ball, carboard tube suspended from the cage lid, nestlet and bedding material. All animals were provided with ad libitum water and standard laboratory chow (Purina, UK) throughout habituation and testing. Details of the sample size age, supplier, history and treatments are summarised in table 1. Sample sizes were based on estimated effects sizes calculated from previous data (Cohen’s d = mean difference/SD) and power calculations. All procedures complied with the UK Animals (Scientific Procedures) Act 1986 and were approved by the University of Bristol Animal Welfare and Ethical Review Body and the UK Home Office.

**Table 1:**
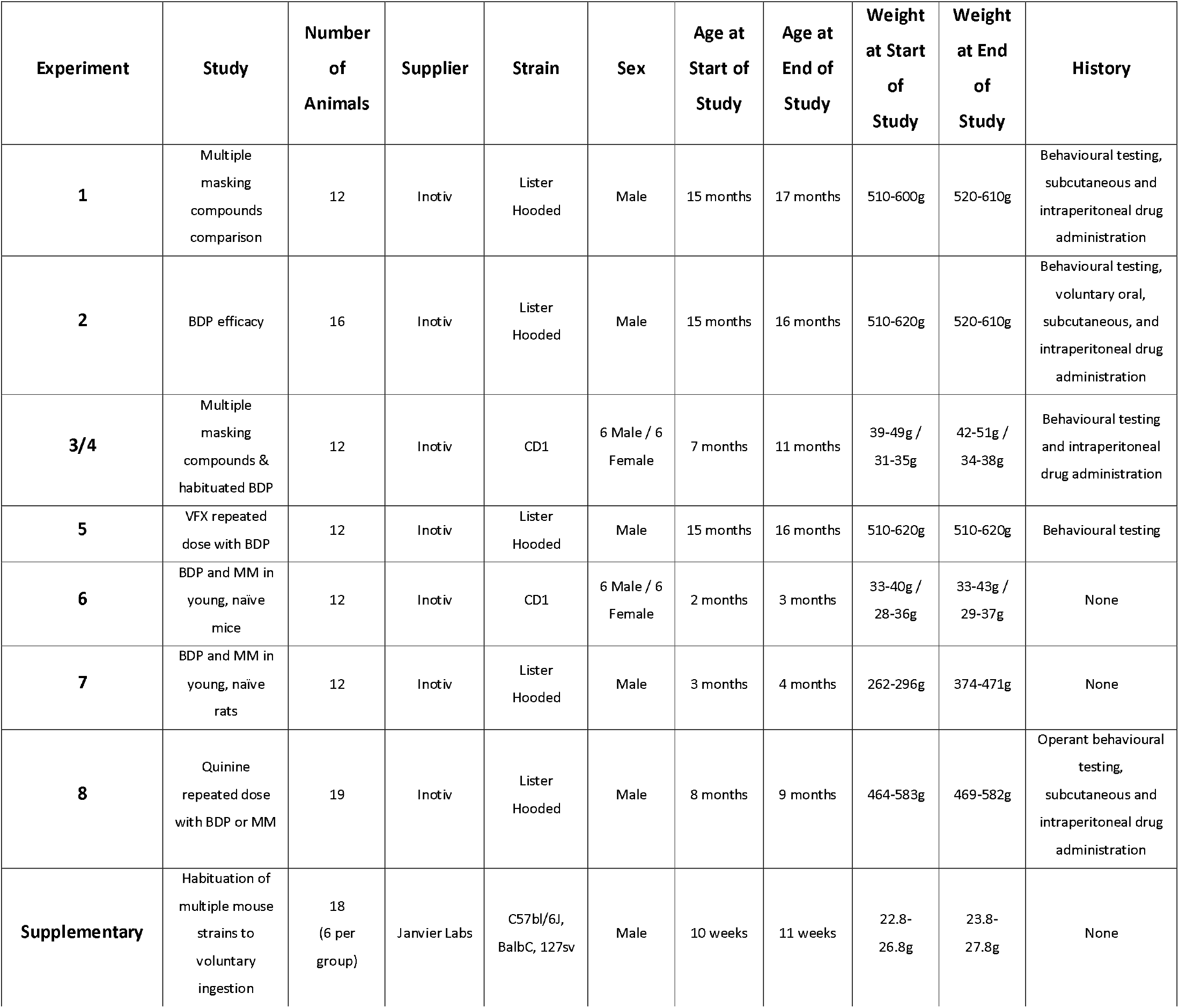
Summary of the different cohorts and treatments used in these studies.

| Experiment | Study | Number of Animals | Supplier | Strain | Sex | Age at Start of Study | Age at End of Study | Weight at Start of Study | Weight at End of Study | History |
| --- | --- | --- | --- | --- | --- | --- | --- | --- | --- | --- |
| 1 | Multiple masking compounds comparison | 12 | Inotiv | Lister Hooded | Male | 15 months | 17 months | 510-600g | 520-610g | Behavioural testing, subcutaneous and intraperitoneal drug administration |
| 2 | BDP efficacy | 16 | Inotiv | Lister Hooded | Male | 15 months | 16 months | 510-620g | 520-610g | Behavioural testing, voluntary oral, subcutaneous, and intraperitoneal drug administration |
| 3/4 | Multiple masking compounds & habituated BDP | 12 | Inotiv | CD1 | 6 Male / 6 Female | 7 months | 11 months | 39-49g / 31-35g | 42-51g / 34-38g | Behavioural testing and intraperitoneal drug administration |
| 5 | VFX repeated dose with BDP | 12 | Inotiv | Lister Hooded | Male | 15 months | 16 months | 510-620g | 510-620g | Behavioural testing |
| 6 | BDP and MM in young, naive mice | 12 | Inotiv | CD1 | 6 Male / 6 Female | 2 months | 3 months | 33-40g / 28-36g | 33-43g / 29-37g | None |
| 7 | BDP and MM in young, naive rats | 12 | Inotiv | Lister Hooded | Male | 3 months | 4 months | 262-296g | 374-471g | None |
| 8 | Quinine repeated dose with BDP or MM | 19 | Inotiv | Lister Hooded | Male | 8 months | 9 months | 464-583g | 469-582g | Operant behavioural testing, subcutaneous and intraperitoneal drug administration |
| Supplementary | Habituation of multiple mouse strains to voluntary ingestion | 18 (6 per group) | Janvier Labs | C57bl/6J, BalbC, 129sv | Male | 10 weeks | 11 weeks | 22.8-26.8g | 23.8-27.8g | None |

### Materials

Animals were dosed with 1mL/kg (rats) or 10mL/kg (mice) of solution using a 1mL luer slip syringe. The palatable vehicle was 10% condensed milk (Carnation) in tap water. Saline (0.9%, Aquapharm®), Saccharine Sodium (0.2%, Sigma UK), Sodium Gluconate (0.3M, ChemCruz Biochemicals), PCCA Bitter Drug Powder (TM) (BDP) (26mg/mL, PCCA UK) and a custom masking mixture (MM) of Saccharine Sodium (Sigma UK), Acesulfame K (fluorochem) and Xanthan Gum (Freee) in equal parts by weight at a concentration of 15mg/mL were tested in combination with the vehicle +/-Quinine Hydrochloride (1mM, Sigma UK) or venlafaxine (3mg/mL Rosemont Pharmaceuticals).

### Habituation

Following arrival in the facility, all animals underwent a handling habituation protocol from day 5 onwards (3Hs-initiative.co.uk) as detailed in table 2.

**Table 2:** Habituation protocols for mice and rats (www.3Hs-initiative.co.uk)

|  |  |
| --- | --- |
| Mouse Habituation | Rat Habituation |
| **Day 1** – open cage and remove all furnishings accept a familiar tube. Keeping the tube within the cage, lift the mouse in the tube and gently encourage to step over your hand back into the cage. Repeat a few times and for each animal in the cage and then place reward into the cage and replace all furniture and lid.<br>**Day 2** – repeat day one but try to have the mouse pause on the hand with gentle cupping each time. With cup handling, the mouse should not feel restrained and should be released if it starts trying to evade handling. For animals which do not require physical handling for the procedures repeat tube handling habituation days 3-5.<br>**Day 3** – for animals requiring physical restraint for procedures progress to cup handling. Repeat day 2 but cup restrain the mouse for a brief period of time before release. Gradually increase the time restrained as the animal's tolerance increases.<br>**Day 4-5** – repeat day 3 increasing time animal is cup restrained.<br>*Individual and strain specific differences will impact on the number of sessions required for the animals to accept each stage and it is important not to progress to a new stage before the animal is comfortable with the current one. | **Day 1** – cage lid off, furniture removed, remaining within the confines of the cage, pick up the rat around the shoulder then immediately put back in cage. Repeat with each cage mate then give reward in home cage.<br>**Day 2** – cage lid off, furniture removed, pick up and transfer to a travel box containing rewards, move all rats from the cage to the travel box, give additional rewards then once consumed, pick up and return to cage, give reward in cage.<br>**Day 3** – open cage lid and wait for rat to approach then pick up, transfer to travel box containing reward, move all rats from the cage to the travel box, give additional rewards then once consumed, pick up and return to cage, give reward in cage.<br>**Day 4/5** – repeat day 3 and start introducing dosing positions and taking palatable solutions from a syringe. If using palatable dose training, this can also serve as the reward.<br>Habituation to sitting on a piece of vet bed can also be useful for certain procedures e.g. subcutaneous injections, administration, or withdrawal of substances through surgically implanted devices. |

Before starting the study, all animals underwent a 1-week habituation to ingesting the palatable vehicle from the syringe following the protocol summarised in figure 1 including decisions to follow to adjust for individual progress. This is the standard amount of time we normally allow for habituation before starting a drug study, but the majority of our animals will consume the entire volume from the syringe on day 2. Data for the habituation of 3 common mouse strains (C57BL6, 127sv, BalbC) is included in the supplementary material (Supplementary figure S1) and shows that there is some variation between strains in drinking time and latency to approach the syringe but, by day 2, all mice are drinking the entire volume.

**Figure 1:**
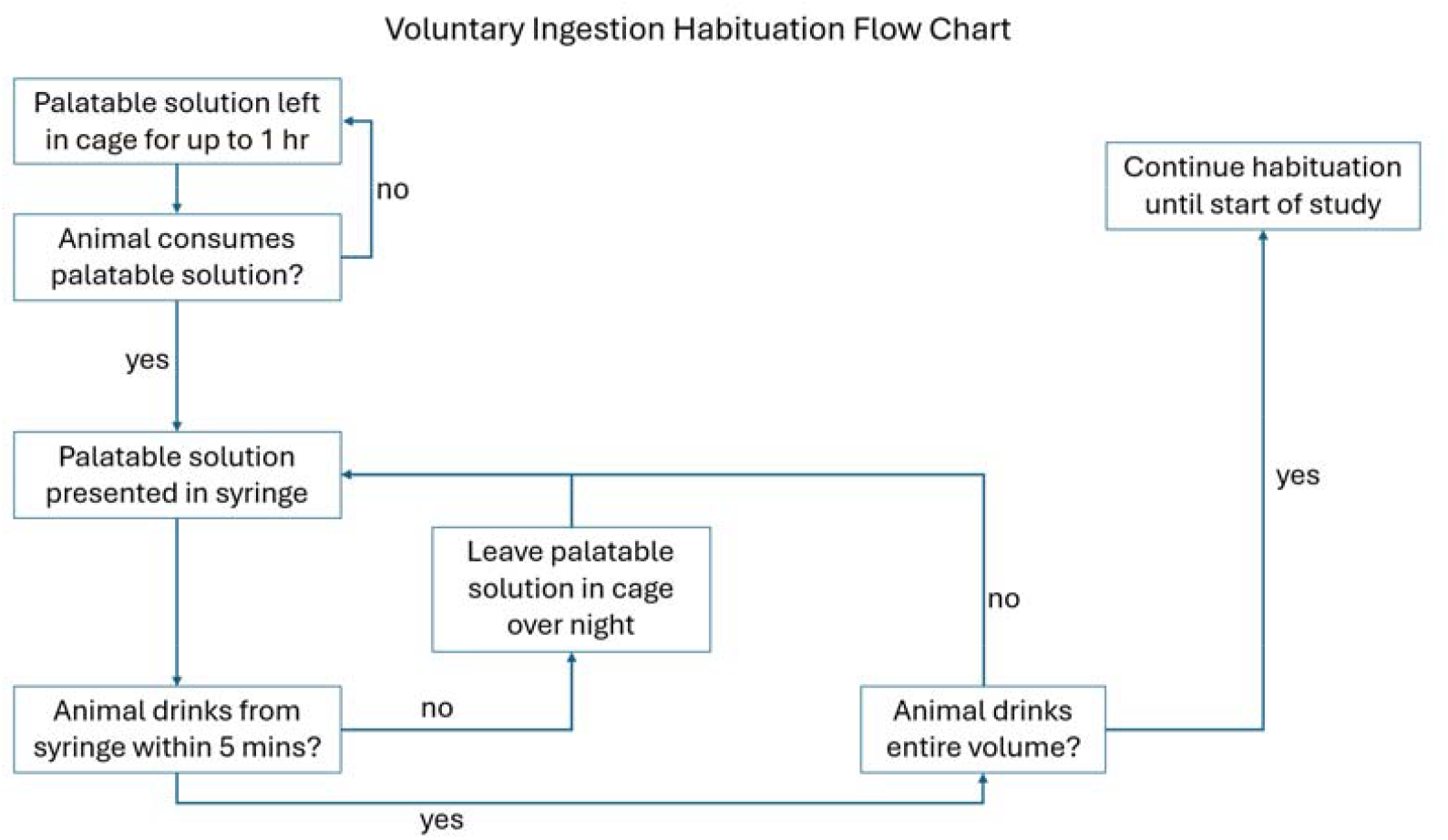
Habituation plan for voluntary ingestion

For rats, all habituation and dosing took place in the animal’s home cage. The animal not being dosed was temporarily removed from the cage for the duration of dosing. The syringe was always presented in the same position through the bars of the cage lid until fully consumed. Rats were alerted to the presence of the syringe by tapping it against the cage lid bars. For group housed mice, animals were moved to a dosing cage identical to their home cage. Mice were alerted by disturbing their food pellets and/or tapping the syringe against the cage lid bars (see supplementary material for dosing videos). Further information on habituation and dosing protocols we used are available from the 3Hs Initiative website (www.3Hs-initiative.com) and are included in the supplementary material.

### Experimental Design

For all experiments, animals received all treatments, including vehicle control, of their study in a fully randomised, within-subject design with the experimenter blind to treatment. For single dose studies treatments were assigned using a counterbalanced system so that the order of treatments was not the same for each animal. For repeated dose studies, animals received the same treatment over 3 consecutive days and received all treatments over the course of the study in a randomised design.

### Experiments 1-4, 6 & 7: Effects of masking compounds on voluntary ingestion of quinine in mice and rats

Each week on day 1-3 and day 5, animals received just the vehicle solution. On day 4 (A.M.), animals received test vehicle-compound combinations and were given up to 5 minutes to consume the entire volume, and on day 4 (P.M.), animals received 0.5 (rats)/0.2mL (mice) vehicle solution (Figure 2). The recorded measures on day 4 were: latency to approach syringe a.m. and p.m., drinking time (max. 5 minutes), and percentage consumed. Latency to approach the syringe was recorded from the initial presentation of the syringe until the first physical contact of the animal with the syringe.

**Figure 2:**
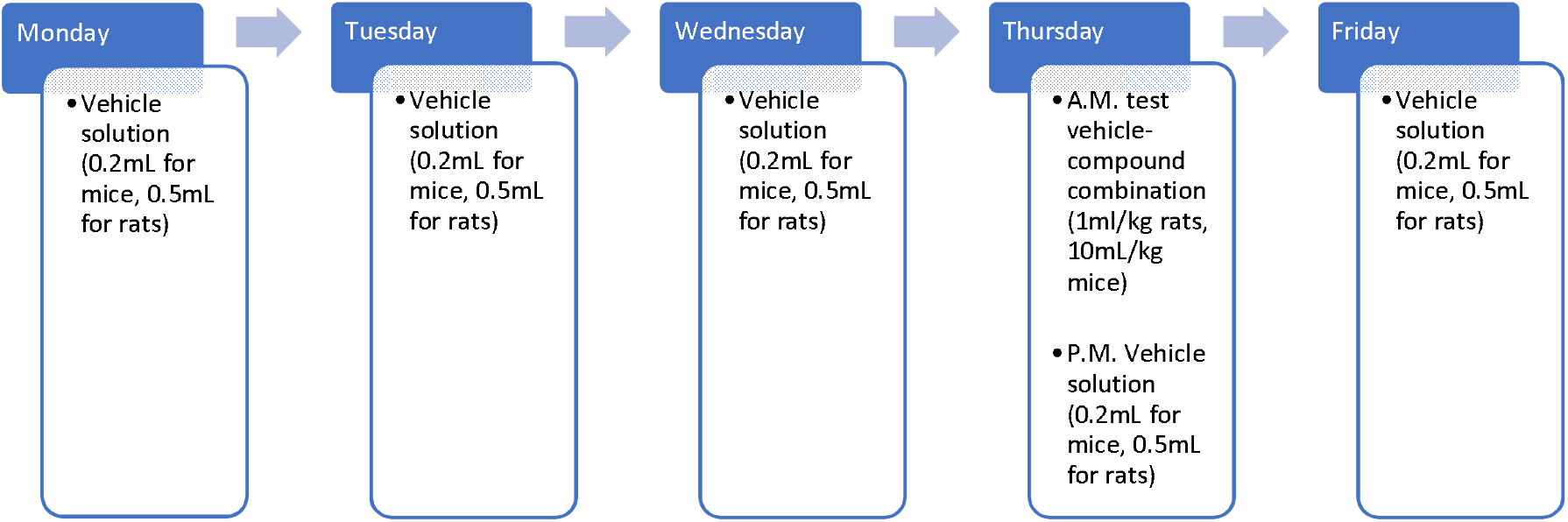
Voluntary ingestion schedule for experiments 1-6

### Experiment 5-8: Effects of BDP and/or MM on repeated dosing with venlafaxine or quinine in rats

On day -1 and 4 pre-habituated animals received vehicle only. On days 1-3, animals received Venlafaxine (3mg/kg) or quinine (1mM) in the palatable vehicle with or without masking agent and were given up to 5 minutes to consume the entire volume (figure 3). The recorded measures for each day were: latency to approach syringe, drinking time (max. 5 minutes), and percentage consumed. Latency to approach the syringe was recorded from the initial presentation of the syringe until the first physical contact of the animal with the syringe.

**Figure 3:**
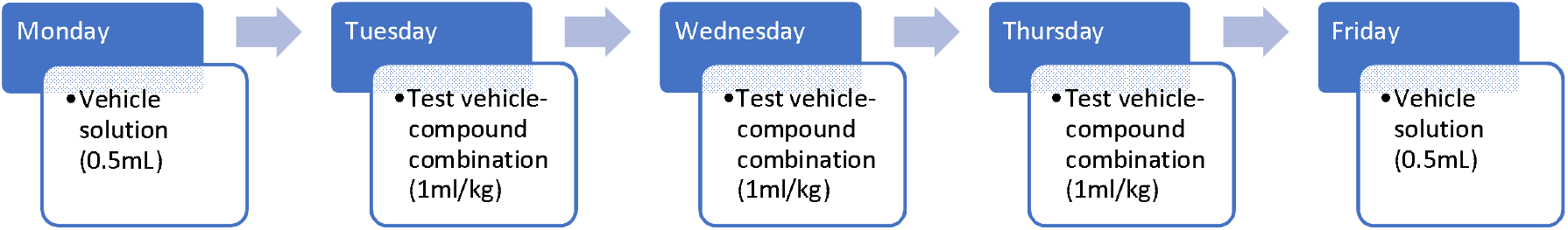
Voluntary ingestion schedule for experiments 7-8

### Statistical Analysis

All data sets were tested for normality using a Shapiro-Wilk test. For experiments 1-4, 5 & 7, where masking agent was the only variable factor, a repeated measures one-way ANOVA with a Geisser-Greenhouse correction was used for normally distributed data sets and a Freidman test was used for non-normally distributed data sets with masking agent as the within-subject factor. Where a significant main effect of treatment was found, post-hoc analysis was conducted using a Holm-Sidak’s multiple comparisons test for normally distributed data sets and a Dunn’s multiple comparisons test for non-normally distributed data sets. For experiments 5 & 8, where both masking agent and day were variable factors, a two-way ANOVA (Treatment and Day) with a Geisser-Greenhouse correction was used and, where a significant effect of masking agent or day was found, post-hoc analysis was conducted using a Holm-Sidak’s multiple comparisons.

## Results

### Experiment 1: Masking compounds commonly used in drug formulations for humans failed to attenuate the effects of quinine on voluntary ingestion (rat)

Reliability of voluntary ingestion was reduced with the addition of quinine which was not improved by the addition of either saline, saccharine or sodium gluconate (Figure 4). There was a main effect of treatment on percentage consumed (F = 14.81, df = 59, p = <0.0001). Quinine decreased consumption (p = 0.0465) but this was not attenuated by quinine-saccharine (p > 0.9999) or quinine-sodium gluconate (p > 0.9999) and was further reduced by quinine-saline (p = 0.0019). There was a main effect of treatment on drinking time (χ^2^ = 32.44, p = <0.0001). Quinine increased drinking time (p = 0.0062) but none of the taste masking compounds attenuated the effect when compared to quinine alone (quinine-saline p = 0.6223, quinine-saccharine p > 0.9999, quinine-sodium gluconate p > 0.9999). There was no effect of treatment on latency to approach for the A.M. or P.M. sessions (A.M. χ^2^ = 2.933 p = 0.569, P.M. χ^2^ = 6.667 p =0.1546).

**Figure 4:**
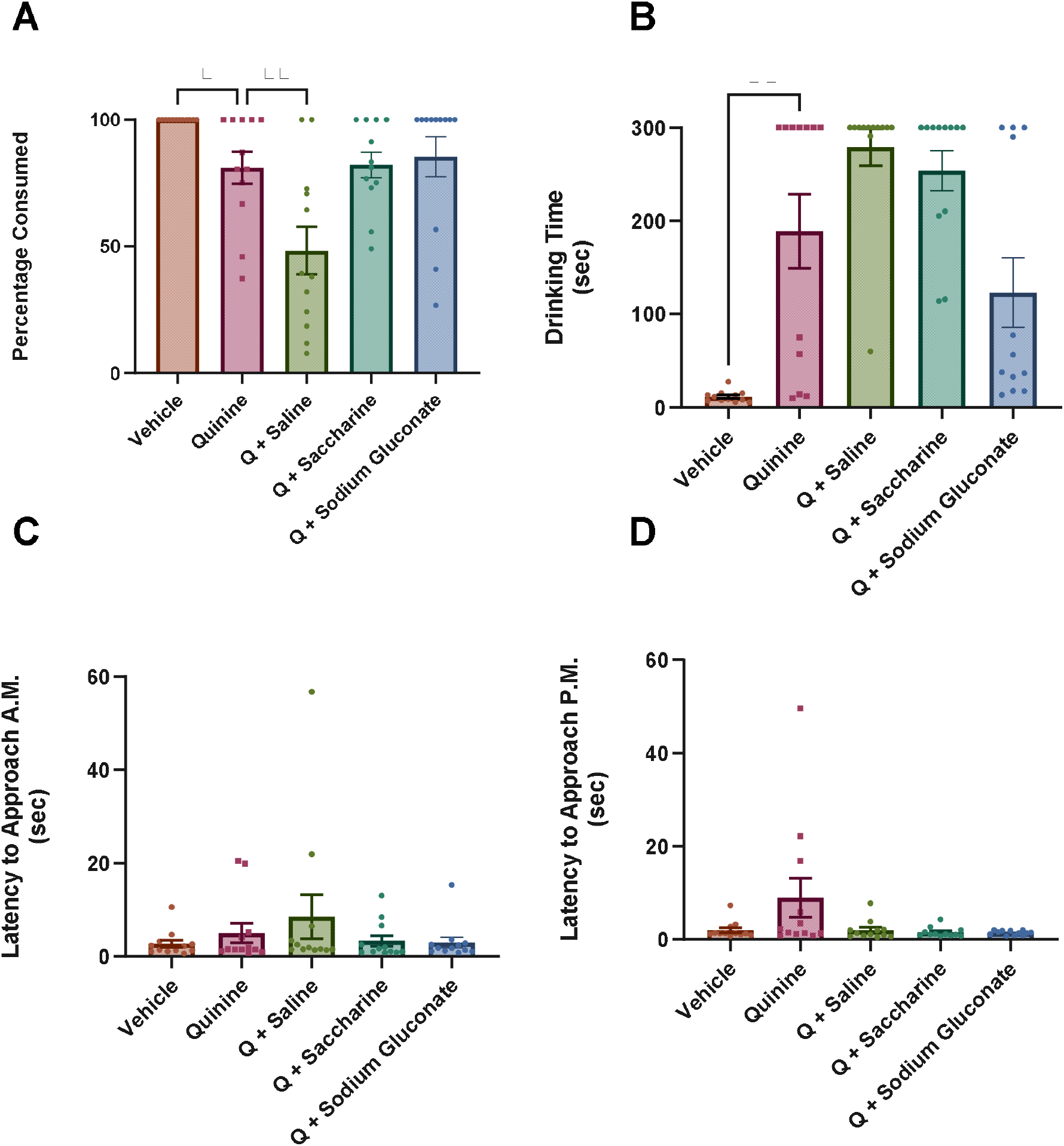
Conventional taste masking compounds had no effect on mitigating the aversive taste of quinine. Rats were presented with vehicle or a quinine solution in 10% condensed milk with or without a taste masking agent. **A** Quinine reduced consumption of the solution (p = 0.0465). None of the taste masking agents attenuated this reduction and saline further decreased the amount consumed (p = 0.0019), **B** Quinine significantly increased drinking time compared to vehicle (p = 0.0062) and the taste masking agents had no significant effect compared to quinine, **C & D** There was no effect on latency to approach the syringe at the A.M. presentation of the test solution or the P.M. presentation of the vehicle. Bars are mean +/-SEM

### Experiment 2: Bitter drug powder effectively masks the bitter taste of quinine and restores voluntary ingestions (rat)

The addition of BDP restored percent consumed and drinking time compared to quinine alone with no difference between vehicle only and quinine-BDP for any of the measures recorded (figure 5). There was a main effect of treatment on the percentage consumed (χ^2^ = 12.00, p = 0.0025). Post hoc analysis showed a reduction in percentage consumed for quinine (p = 0.0031) compared to vehicle which was attenuated by addition of the BDP (p = 0.0031 vs quinine). There was a main effect of treatment on drinking time (χ^2^ = 17.38, p = 0.0002). Quinine increased drinking time compared to vehicle (p < 0.0001) and quinine-BDP reduced drinking time to vehicle levels (p = 0.0094 vs quinine). There was no effect of treatment on latency to approach for the A.M. or P.M. sessions (A.M. χ^2^ = 1.625 p = 0.4437, P.M. χ^2^ = 2.0 p =0.3679).

**Figure 5:**
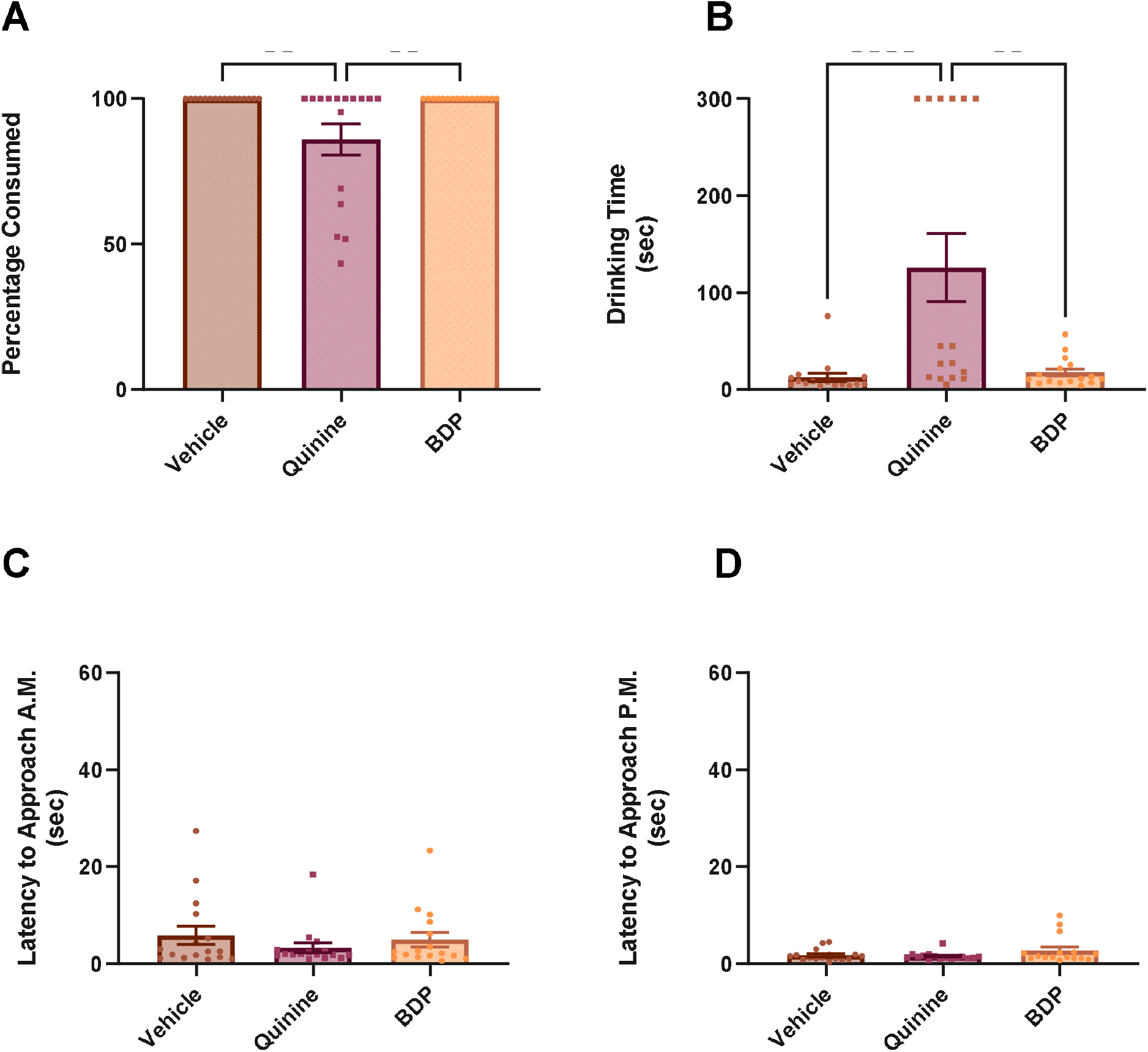
Bitter drug powder^tm^ (BDP) was effective at masking the aversive taste of quinine. Rats were presented with vehicle or a quinine solution in 10% condensed milk with or without BDP. **A** Quinine significantly decreased the amount consumed (p < 0.0001) and the addition of BDP significantly increased consumption compared to quinine (p = 0.0094), **B** Quinine significantly increased drinking time compared to vehicle (p = 0.0031) and BDP significantly reduced drinking time compared to quinine (p = 0.0031), **C & D** There was no effect on latency to approach the syringe at the A.M. presentation of the test solution or the P.M. presentation of the vehicle. Bars are mean +/-SEM.

### Experiments 3 & 4: Bitter Drug Powder effectively masks the bitter taste of quinine and improves voluntary ingestion (mouse)

Mice reduced their consumption and increased drinking time when administered quinine in the palatable vehicle. Of the masking agents tested, only BDP was effective at masking these effects with no effects observed for saline, saccharine or sodium gluconate when compared to quinine alone (figure 6). The percent consumed and drinking time for the quinine-BDP combination was not different from vehicle alone although full reversal of the effects were not observed for all animals. There was a main effect of treatment on the percentage consumed (χ^2^ = 33.34, p = <0.0001). Post hoc analysis showed a reduction in percentage consumed for quinine (p = 0.0024). Quinine-BDP increased percentage consumed compared to quinine (p = 0.376) and there was no significant difference in percentage consumed for quinine-saline (p > 0.9999) quinine-saccharine (p > 0.9999) or quinine-sodium gluconate (p = 0.9522) compared to quinine. There was a main effect of treatment on drinking time (χ^2^ = 39.53, p = <0.0001). Quinine increased drinking time compared to vehicle (p = 0.0001) and quinine-BDP reduced drinking time compared to quinine (p = 0.319). None of the other masking compounds attenuated the effect on drinking time when compared to quinine (quinine-saline p > 0.9999, quinine-saccharine p > 0.9999, quinine-sodium gluconate p > 0.9999). There was no effect of treatment on latency to approach for the A.M. or P.M. sessions (A.M. χ^2^ = 9.286 p = 0.0982, P.M. χ^2^ = 4.81 p = 0.4396).

**Figure 6:**
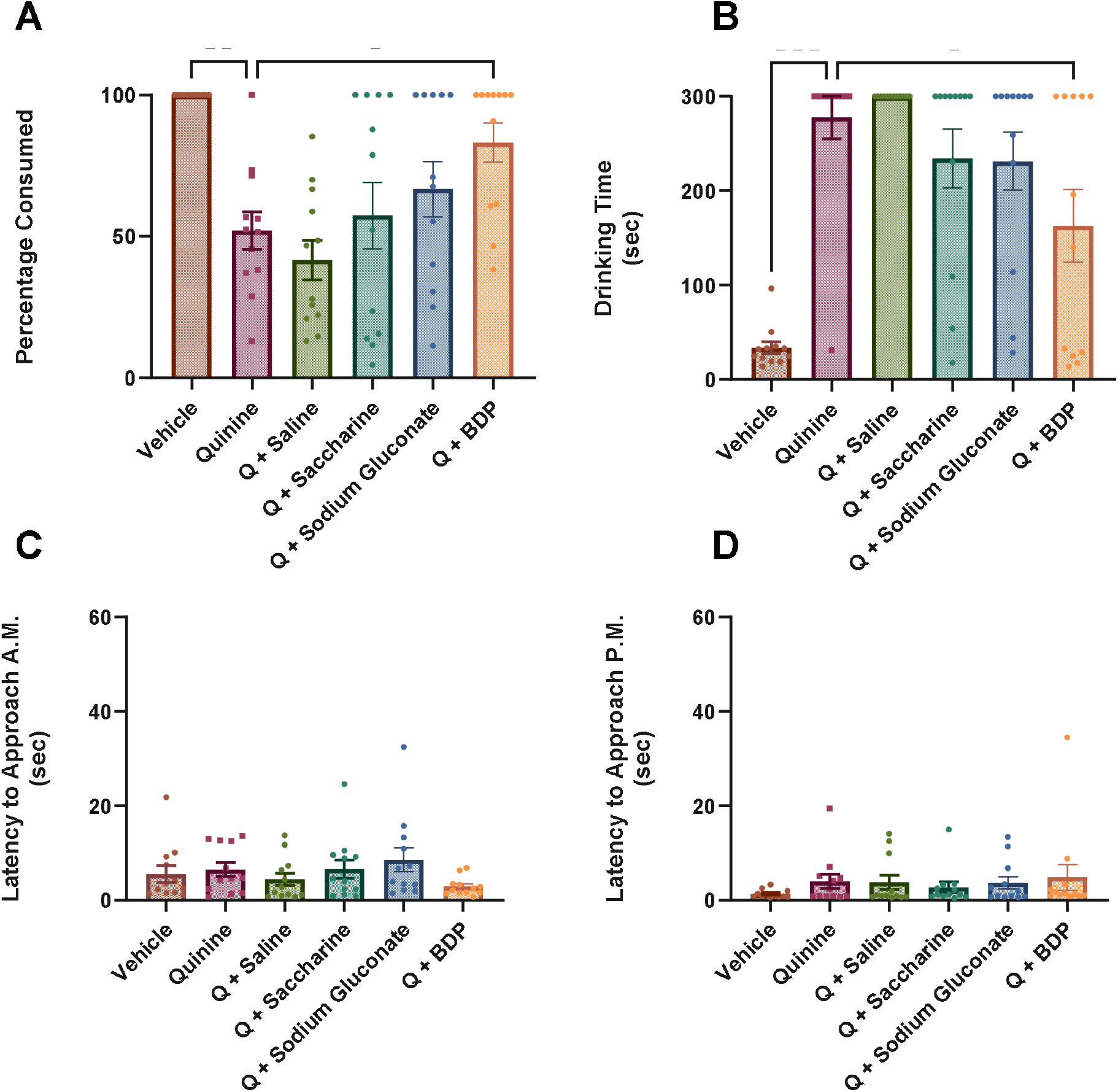
Bitter drug powder^tm^ (BDP) but no conventional taste masking agents were effective at masking the aversive taste of quinine. Mice were presented with vehicle or a quinine solution in 10% condensed milk with or without BDP. **A** Quinine significantly decreased the amount consumed (p = 0.0001) and the addition of BDP significantly increased consumption compared to quinine (p = 0.0319), **B** Quinine significantly increased drinking time compared to vehicle (p = 0.0024) and BDP significantly reduced drinking time compared to quinine (p = 0.0376), **C & D** There was no effect on latency to approach the syringe at the A.M. presentation of the test solution or the P.M. presentation of the vehicle. Bars are mean +/-SEM

Although the mice were habituated to the 10% condensed milk prior to the study, the BDP changes the consistency of the solution which may have contributed to BDP failing to fully restore consumption. We therefore used an additional 5-day habituation to the vehicle-BDP combination before repeating the test with quinine. BDP fully reversed the effects of quinine in all animals in terms of percent consumed and drinking time (figure 7). There was a main effect of treatment on the percentage consumed (χ^2^ = 13.85, p = 0.001). Post hoc analysis showed a reduction in percentage consumed for quinine (p = 0.0025) compared to vehicle which was fully attenuated by addition of the BDP (p = 0.0025 vs quinine). There was a main effect of treatment on drinking time (χ^2^ = 16.17, p = 0.0003). Quinine increased drinking time compared to vehicle (p = 0.0002) and quinine-BDP reduced drinking time to vehicle levels (p = 0.0085 vs quinine). There was no effect of treatment on latency to approach for the A.M. or P.M. sessions (A.M. χ^2^ = 1.167 p = 0.558, P.M. χ^2^ = 2.596 p = 0.2731).

**Figure 7:**
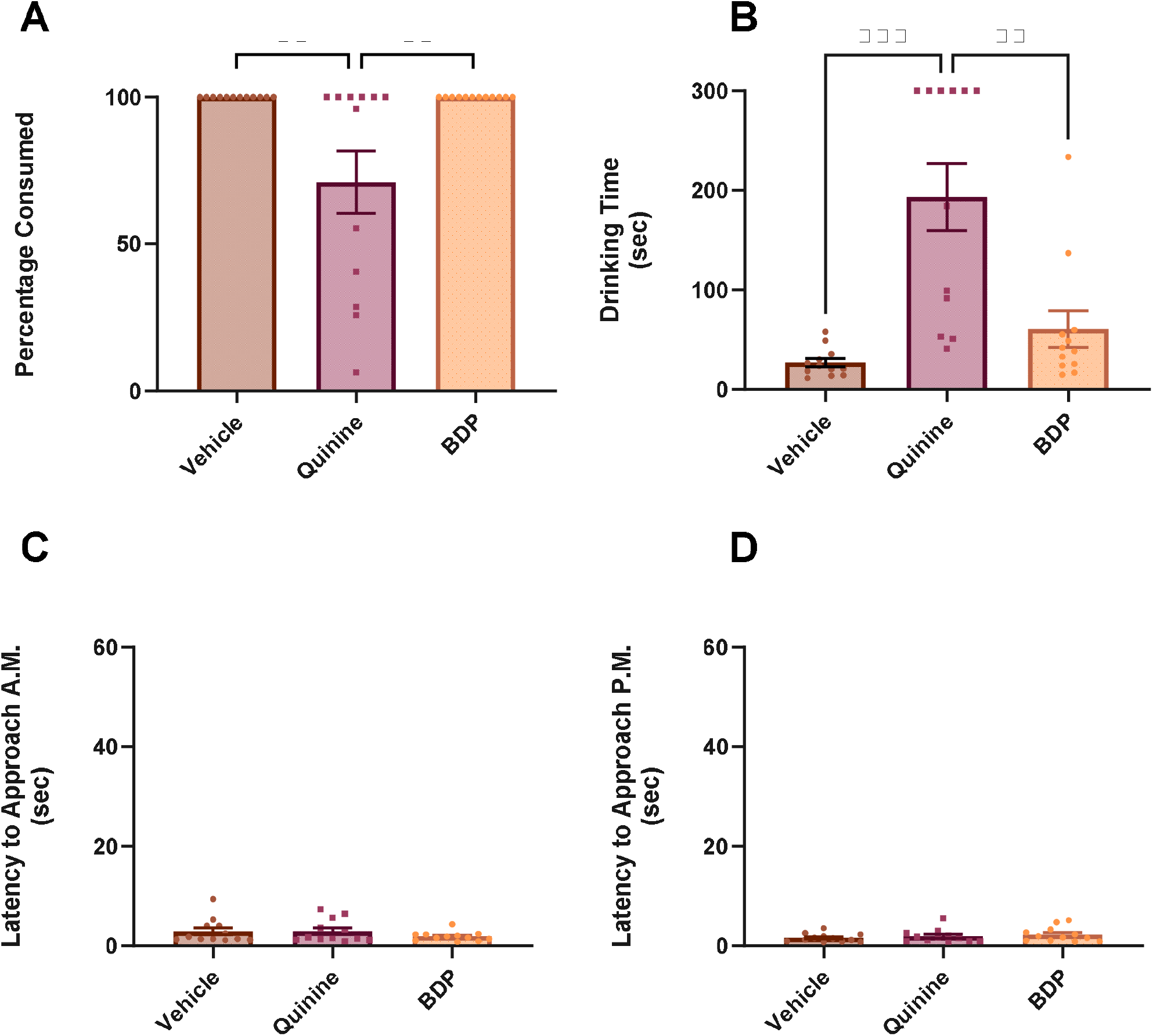
Bitter drug powder^tm^ (BDP) was effective at masking the aversive taste of quinine. Mice were presented with vehicle or a quinine solution in 10% condensed milk with or without BDP. **A** Quinine significantly decreased the amount consumed and the addition of BDP significantly increased consumption compared to quinine (p = 0.0025), **B** Quinine significantly increased drinking time compared to vehicle (p = 0.0002) and BDP significantly reduced drinking time compared to quinine (p = 0.0085), **C & D** There was no effect on latency to approach the syringe at the A.M. presentation of the test solution or the P.M. presentation of the vehicle. Bars are mean +/-SEM

### Experiment 5 Venlafaxine, repeated dose +/-BDP (rat)

Rats reduced their intake of venlafaxine (VFX) with repeated administration, but this effect was fully prevented by the addition of BDP (figure 8). There was a main effect of treatment (F _(1,22)_ = 18.68, p = 0.0003) and day (F _(1.827, 40.18)_ = 7.962, p = 0.0016) on percentage consumed and an interaction between treatment and day (F _(3, 66)_ = 7.723, p = 0.0002). Percentage consumed was reduced in the of Vehicle+VFX group on days 2 and 3 compared to day -1 (day 1 p = 0.4141 p = day 2 p = 0.0307, day 3 p = 0.0077). This was attenuated by addition of BDP with percentage consumed of BDP+VFX increased on day 3 (p = 0.0117) compared to VFX alone. There was a main effect of treatment (F _(1,22)_ = 22.01, p = 0.0001) and day (F _(2.459, 54.10)_ = 15.13, p = <0.0001) on drinking time and an interaction between treatment and day (F _(2.459, 54.10)_ = 11.63, p = <0.0001). VFX increased drinking time with repeated administration which was attenuated by the addition of the BDP. Drinking time of Vehicle+VFX increased on days 2 and 3 compared to day -1 (day 1 p = 0.2215, day 2 p = 0.002, day 3 p = 0.0002) and drinking time of BDP+Vehicle was decreased on days 2 and 3 (day 2 p = 0.0131, day 3 p = 0.0003) compared to VFX alone. There was a main effect of treatment on latency to approach (F _(1,22)_ = 4.740, p = 0.0405) but no significant post hoc differences.

**Figure 8:**
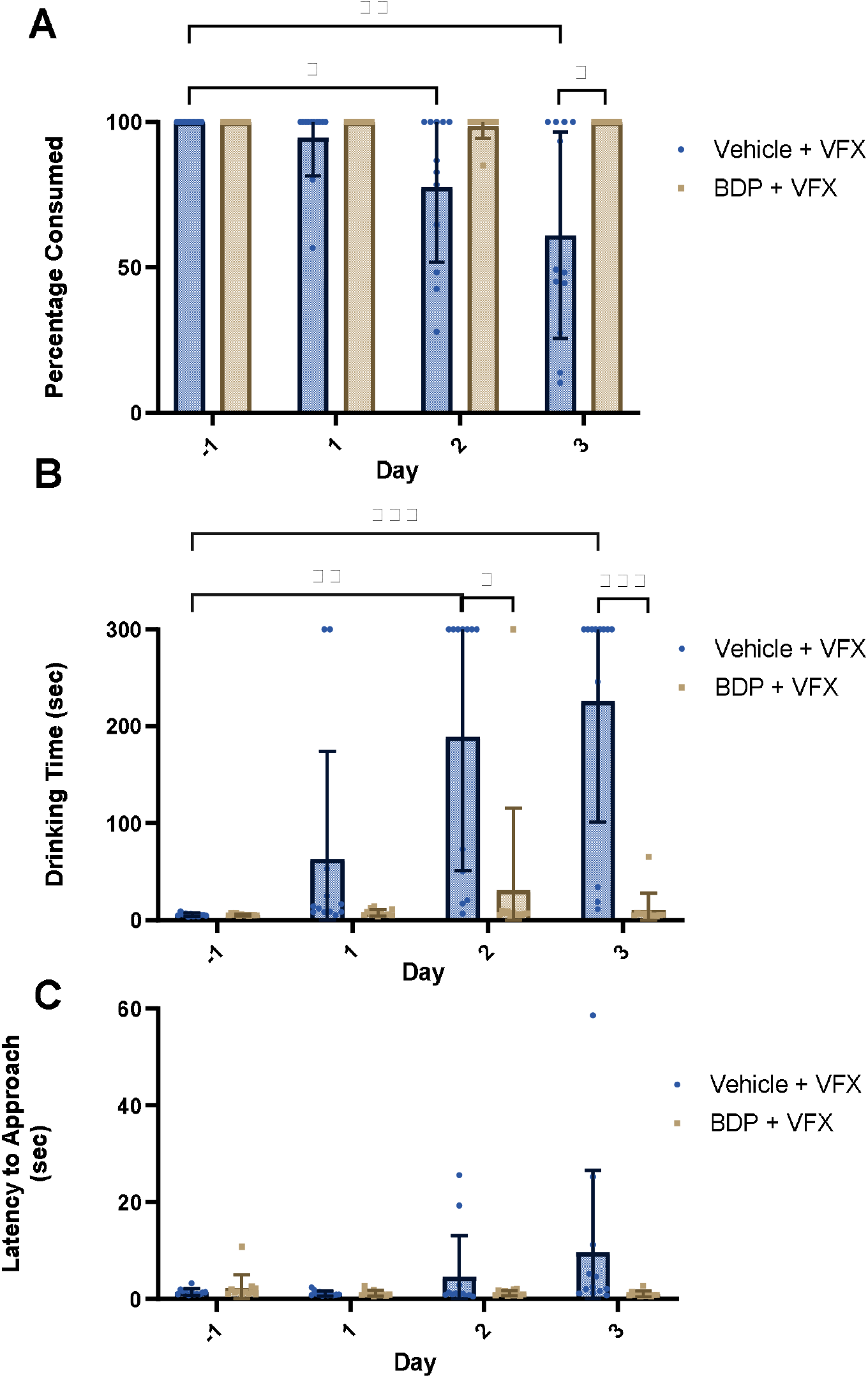
Bitter drug powder^tm^ (BDP) was effective at masking the aversive taste of Venlafaxine (VFX) over successive dosing days. Rats were presented with VFX in 10% condensed milk with or without BDP. **A** Vehicle + VFX significantly reduced amount consumed on days 2 and 3 compared to day -1 (p = 0.0307, p = 0.0077) BDP significantly increased the amount consumed (p = 0.0117) on day 3 of dosing, **B** Vehicle + VFX significantly increased drinking time on days 2 and 3 compared to day -1 (p = 0.002, p = 0.0002) BDP significantly decreased drinking time on days 2 (p = 0.0131) and 3 (p = 0003) of dosing, **C** There was no effect on latency to approach the syringe at the A.M. presentation of the test solution. Bars are mean +/-SEM.

### Experiments 6 & 7: Masking Mixture effectively masks the bitter taste of quinine and restores voluntary ingestions (rat and mouse)

The combination of the two artificial sweeteners and xanthan gum, in our own formulation of a masking mixture, was as effective as BDP in masking the effects of quinine restoring voluntary ingestion to vehicle levels in rats and mice. In the rat study (figure 9), there was a main effect of treatment on the percentage consumed (χ^2^ = 21.00, p = 0.0001). Post hoc analysis showed a reduction in percentage consumed for quinine (p = 0.0269) compared to vehicle, BDP and MM. Percentage consumed for both MM and BDP remained at 100%. There was a main effect of treatment on drinking time (χ^2^ = 22.0, p < 0.0001). Quinine increased drinking time compared to vehicle (p = 0.0002) and quinine-MM and quinine-BDP attenuated this to vehicle levels (MM p = 0.003, BDP p = 0.0009 vs quinine). There was no effect of quinine +/-masking compound on latency to approach for the A.M. or P.M. sessions (A.M. χ^2^ = 6.9 p = 0.0752, P.M. χ^2^ = 4.5 p =0.2123).

**Figure 9:**
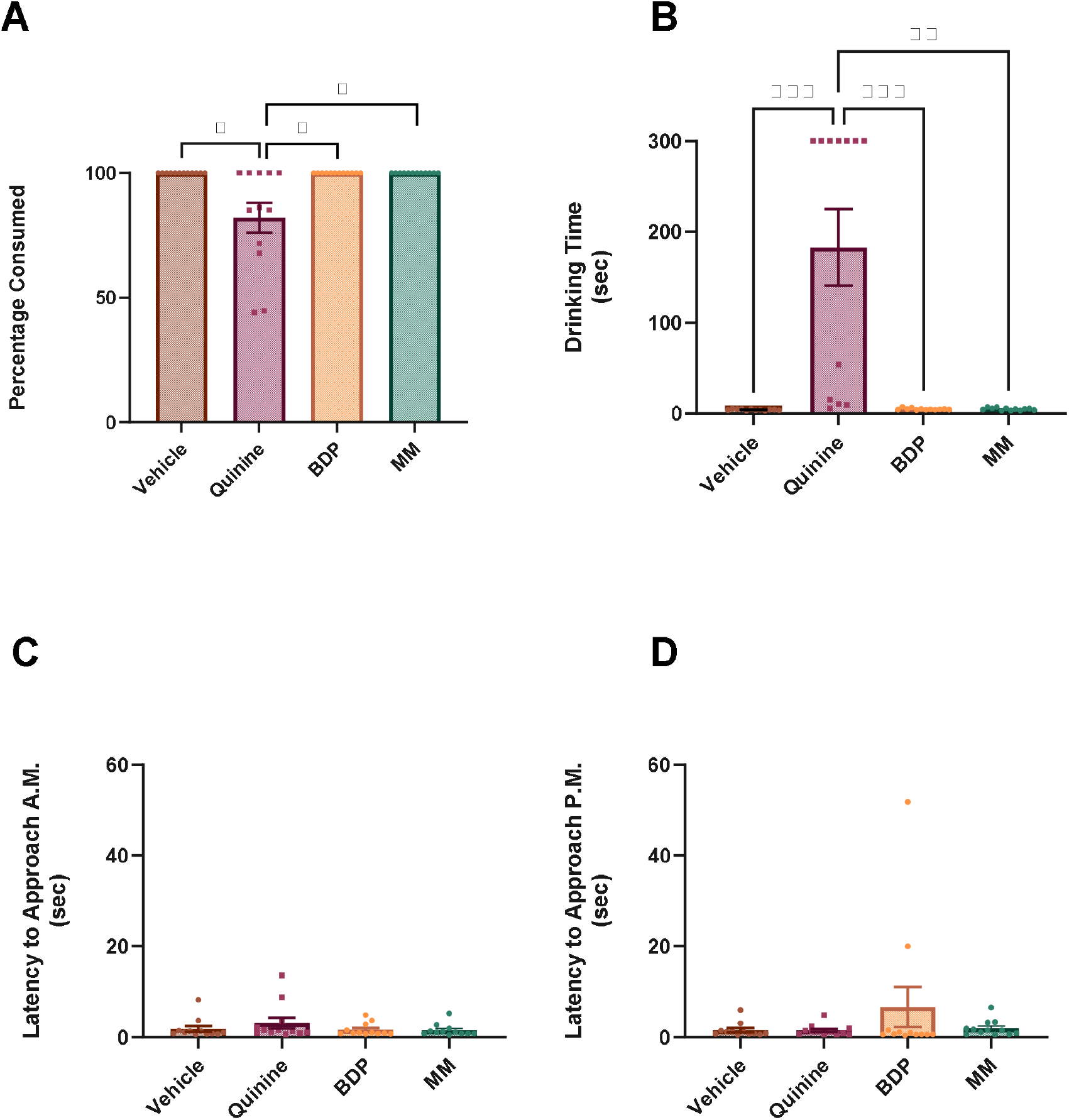
Bitter drug powder^tm^ (BDP) and masking mixture (MM) were effective at masking the aversive taste of quinine. Rats were presented with vehicle or a quinine solution in 10% condensed milk with or without BDP or MM. **A** Quinine significantly decreased the amount consumed compared to vehicle, BDP, and MM (p = 0.0269), **B** Quinine significantly increased drinking time compared to vehicle (p = 0.0002) and both BDP and MM significantly reduced drinking time compared to quinine (p = 0.0009, p = 0.003), **C & D** There was no effect on latency to approach the syringe at the A.M. presentation of the test solution or the P.M. presentation of the vehicle. Bars are mean +/-SEM.

In mice (figure 10), there was a main effect of treatment on the percentage consumed (χ^2^ = 27.00, p < 0.0001). Post hoc analysis showed a reduction in percentage consumed for quinine (p = 0.0266) compared to vehicle. Percentage consumed for both MM and BDP remained at 100%. There was a main effect of treatment on drinking time (χ^2^ = 22.3, p < 0.0001). Quinine increased drinking time compared to vehicle (p < 0.0001) and quinine-MM and quinine-BDP attenuated this to vehicle levels (MM p = 0.016, BDP p = 0.0094 vs quinine). There was no effect of treatment on latency to approach for the A.M. or P.M. sessions (A.M. χ^2^ = 5.8 p = 0.1218, P.M. χ^2^ = 3.1 p =0.3765).

**Figure 10:**
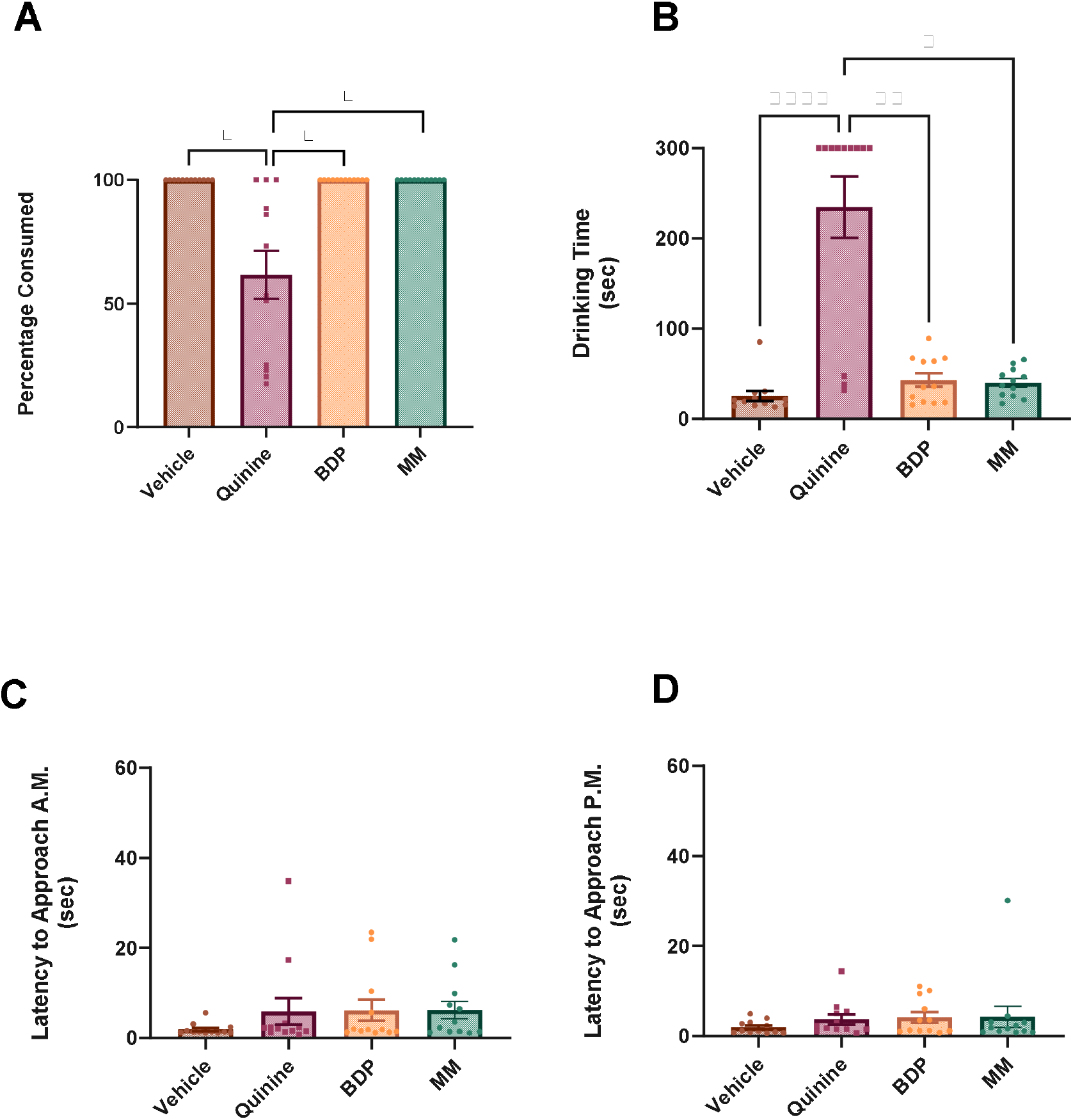
Bitter drug powder^tm^ (BDP) and masking mixture (MM) were effective at masking the aversive taste of quinine. Mice were presented with vehicle or a quinine solution in 10% condensed milk with or without BDP or MM. **A** Quinine significantly decreased the amount consumed compared to vehicle, BDP, and MM (p = 0.0266), **B** Quinine significantly increased drinking time compared to vehicle (p < 0.0001) and both BDP and MM significantly reduced drinking time compared to quinine (p = 0.0094, p = 0.016), **C & D** There was no effect on latency to approach the syringe at the A.M. presentation of the test solution or the P.M. presentation of the vehicle. Bars are mean +/-SEM

### Experiment 8 Quinine, repeated dose +/-MM or BDP (rat)

In this study, we used a within-subject design to compare the efficacy of MM and BDP on quinine voluntary-ingestion over 3 consecutive days of dosing. Similar to the findings with VFX, animals consumed the quinine on the first session, but this declined with repeated administration, an effect which was prevented by both BDP and MM (figure 11). There was a main effect of treatment (F _(2, 54)_= 30.09, p < 0.0001) and day (F _(2.484, 134.1)_ = 14.62, p < 0.0001) on percentage consumed and an interaction between treatment and day (F _(4.968, 134.1)_ = 14.62, p = <0.0001). Percentage consumed was reduced on days 1, 2 and 3 compared to day -1 (day 1 p = 0.0031 p = day 2 p = 0.0002, day 3 p = 0.0002). This was attenuated by both MM and BDP (day 1 p = 0.0133, day 2 p = 0.001, day 3 p = 0.0009). There was a main effect of treatment (F _(2, 54)_ = 45.89, p < 0.0001) on drinking time and an interaction between treatment and day (F _(4.905, 132.4)_ = 23.64, p = <0.0001). Quinine alone resulted in a decrease in consumption and increased drinking time with repeated administration on all days (day 1, 2 and 3 p < 0.0001) which was attenuated by the addition of MM and BDP (day 1 MM p < 0.0001 BDP p = 0.0001, day 2 MM p < 0.0001 BDP p < 0.0001, day 3 MM p = 0.0002 BDP p = 0.0002). There was a main effect of treatment on latency to approach (F _(2, 54)_ = 3.286, p = 0.045) but no significant post hoc results.

**Figure 11:**
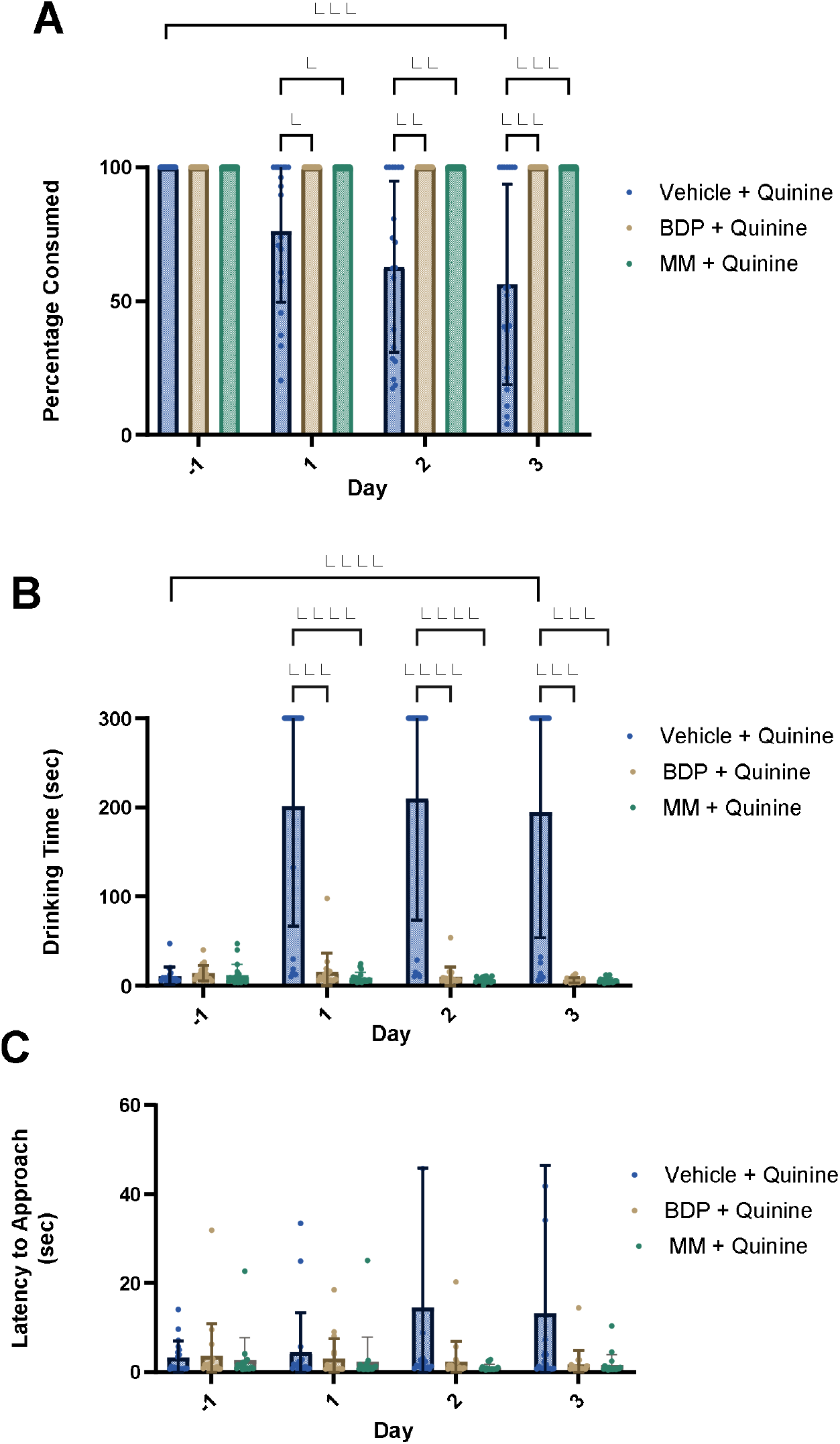
Bitter drug powder^tm^ (BDP) and masking mixture (MM) were effective at masking the aversive taste of Venlafaxine (VFX) over successive dosing days. Rats were presented with VFX in 10% condensed milk with or without BDP or MM. **A** Vehicle + VFX significantly reduced amount consumed on all days compared to both BDP and MM (day 1 p = 0.0133, day 2 p = 0.001, day 3 p = 0.0009) and percentage consumed of quinine was reduced on days 2 and 3 compared to day -1 (day 1 p = 0.0031 p = day 2 p = 0.0002, day 3 p = 0.0002) the addition of BDP or MM resulted in a maintenance of the 100% consumption rate for all animals, **B** BDP and MM significantly decreased drinking time on all days (day 1 p = 0.0001/<0.0001, day 2 p < 0.0001/<0.0001, day 3 p = 0.0002/0.0002, **C** There was no effect on latency to approach the syringe at the A.M. presentation of the test solution. Bars are mean +/-SEM.

## Discussion

In both rats and mice, the addition of quinine to the palatable vehicle reduced the percentage consumed and increased drinking time, demonstrating that quinine’s aversive taste reduces the reliability of voluntary ingestion. These effects were more marked in mice than in rats possibly due to the larger dose volume used in mice and longer drinking times. In rats and mice, we found no beneficial effects with the addition of saline, saccharine or sodium gluconate contrary to the effects seen in human studies (28,29,31). Although these compounds have a masking effect sufficient in humans, they do not sufficiently mask the bitter taste of quinine to achieve reliable ingestion in rats and mice. Interestingly, in rats, the addition of saline had a negative impact on consumption possibly due to species differences in the expression of taste receptors which may result in rodents being more sensitive to certain tastes than humans (35). In contrast, the Bitter Drug Powder™ was highly effective in masking the bitter taste of the quinine solution in both rats and mice improving reliability of voluntary ingestion in terms of both percentage consumed and reduced drinking time. In rats we saw a return to 100% ingestion and similar consumption time to vehicle alone. In mice, the Bitter Drug Powder™ significantly improved the percentage consumed and the consumption rate compared to quinine but, in the initial study, did not result in 100% consumption. We hypothesised that this could be because the addition of the Bitter Drug Powder™ resulted in a more viscous solution than vehicle alone which, while having no impact on ingestion in the rats, resulted in a neophobic reaction in the mice (23). Prior habituation to the more viscous solution resulted in 100% consumption of the quinine-BDP solution and comparable drinking time to vehicle alone. This highlights the need to consider all aspects of the vehicle solution, not just the taste but also the texture. Although we found no beneficial effects on ingestion with saline, saccharin or sodium-gluconate in mice or rats, these findings with BDP suggest further studies which include additional habituation sessions to each of these vehicles may improve their efficacy.

To further evaluate the efficacy of BDP in the context of a drug study, we tested rats using repeated dosing with the serotonin and noradrenaline re-uptake inhibitor and antidepressant, venlafaxine. We have previously tested this drug using the Affective Bias Test (ABT) via intraperitoneal injection and by voluntary ingestion showing comparable efficacy (39,40) but have seen a progressive decrease in the willingness of rats to consume the dose over consecutive dosing days. This effect was similarly observed in the current study but was fully mitigated using BDP demonstrating the benefits of using this masking agent to improve the reliability of voluntary ingestion in a pharmacological experiment where previous issues had limited its implementation.

One potential issue with integrating BDP into scientific studies is its proprietary formulation with the exact details of the composition unknown. To support future use of masking agents for scientific studies we developed our own formulation using a mixture of artificial sweeteners and xanthan gum and tested using quinine in a within-subject design (mice and rats) using repeated dosing. MM fully attenuated the effects of quinine on consumption and drinking time in both rats and mice with equivalent efficacy to BDP. Using a repeated dosing study design with quinine in rats, we observed the same progressive decrease in consumption that was fully mitigated by the addition of either MM or BDP. Rats maintained a 100% consumption rate over the 3 days, and their drinking time remained short. This also reassured us that the similar decrease in consumption of both VFX and quinine administered over consecutive days was likely due to the taste of the solution and not a conditioned aversion to the effects of venlafaxine, which we would not expect to be mitigated so effectively by a taste masking compound. Taken together, these findings demonstrate that MM is effective at masking bitter taste and, as the exact composition of this mixture is known, it can be integrated in scientific studies where the exact vehicle composition is important. This also allows us to ensure that the masking formulation does not contain anything that might interact with the drugs being administered.

The repeated dosing studies were interesting in terms of a comparison with the previous within-subject design experiments where we observed that some animals consumed 100% of the quinine despite its bitter taste. Integrating these results with the findings from the repeated dosing with both venlafaxine and quinine suggests some animals, particularly rats, will initially consume a bitter drug in the palatable solution but then show reduced willingness to voluntarily-ingest with repeated administration. This limits the reliability of voluntary-ingestion and further supports the benefits of including a masking agent in studies where repeated administration is needed.

In all studies we saw no effect on the latency to approach the syringe during the A.M. or P.M. sessions for any of the vehicle or vehicle-test compound solutions. This shows that the inhibition of drinking was unlikely to be due to a conditioned aversion caused by negative side effects of the test solutions. Rodents have been shown to associate negative drug effects with the delivery vehicle leading to a reduction in willingness to interact with or consume the vehicle even without the drug (37,38). This would predict this would cause an increase in latency to approach in the P.M. session following the aversive experience even when presented with just the vehicle solution.

To reduce the numbers of animals used for regulated procedures, we used mice and rats which had been involved in other regulated procedures however, their prior experience may have influenced their compliance with voluntary ingestion therefore we also included experiments in young, treatment-naïve rats and mice (experiments 5, 6 and 8). We found similar results in these animals demonstrating voluntary ingestion can be used reliably in animals that had not undergone extensive prior handling, vehicle habituation or drug testing. This increased our confidence that taste masking compounds, combined with our habituation protocols, are effective across a range of ages regardless of prior experience.

The principles of the 3Rs and the legal frameworks that we work within dictate that, when designing studies, the most refined methods suitable for the programme of work should be used (41,42). Oral gavage is the most frequently used route of administration (based on data from past 5 years, supplementary table S1) and the route most often used in patients. Voluntary ingestion offers a significant refinement over oral gavage by eliminating the risk of injury to the animal and the distress of the procedure so should be considered for all oral dosing studies. Currently, the reluctance to adopt voluntary ingestion as the preferred oral dosing method and the justification for gavage may be due to its perceived lack of reliability. The findings from this study demonstrate that the addition of a taste masking agent (BDP or MM) to the drug formulation improves the reliability of voluntary ingestion as a method for both acute and repeat-dosing studies. Methods that involve placing the test compound in a palatable food in the animal’s home cage, or mixing it with their food or water, introduces uncertainty in terms of the actual dose consumed and the precise time of dosing. While this is may be acceptable for some studies or veterinary treatments, many studies require greater control over these factors. Using syringe/pipette dosing, where the volume administered can be exactly measured and consumption is observed by the experimenter, eliminates uncertainty in terms of volume, time, and accuracy of dosing. A potential barrier to adoption of this refined method is the time required habituate the animals and time taken to administer the dose. The habituation protocol (see supplementary material) takes between 3 and 5 days and can be done during the animals’ acclimatisation period; therefore, does not need to delay the experimental programme. The dosing times we recorded in this study showed that average drinking time was ∼40 seconds for a mouse and ∼14 seconds for a rat. There is also the possibility that one person can dose two animals simultaneously using this method making the time requirement of voluntary ingestion at least equivalent to oral gavage if not quicker.

In these studies, we used a 10% condensed milk solution as the vehicle. This does present the potential for an effect of both the sugar content and milk proteins however this can be managed by using different palatable vehicles. The benefit of taste masking compounds is that they are compatible with a variety of vehicles, providing different options depending on the study objectives. We have previously found that rodents will voluntarily ingest a variety of palatable solutions from a syringe e.g. fruit juice, nut milk, peanut oil, and combining these with taste masking compounds should provide researchers with a number of options compatible with the study objectives. We have focused on water soluble compounds as these are easy to formulate into a suitable solution for oral ingestion. This might be more complicated for drugs that are water-insoluble, and more work is needed to determine the most appropriate ways to prepare these compounds in a way that is compatible with voluntary ingestion. This could involve using low concentrations of solvents like ethanol or dimethyl sulphoxide in combination with palatable solutions or using oil-based vehicles like peanut butter/peanut oil. The taste masking compounds that we found to be effective also increase the viscosity of the vehicle so it may also be possible to dose water-insoluble drugs as a suspension, but this would require further study and validation.

## Conclusion

We investigated whether compounds conventionally used for taste masking drugs in humans could be effective at improving the reliability of our voluntary oral ingestion protocol in rats and mice. We showed that several compounds that are effective in human trials were ineffective when used for rats and mice. However, both Bitter Drug Powder™ and our own Masking Mixture were highly effective at masking the aversive taste of quinine and venlafaxine, a drug that had been previously shown to be aversive and maintained 100% consumption rate and short drinking time in both the within-subject design and repeated administration over 3 consecutive days. These findings provide evidence to support an effective taste masking protocol that should enable a broader range of drugs to be administered by voluntary ingestion, thus reducing the need for oral gavage and achieving a refinement.

## Acknowledgements

Funding for this research was provided by a BBSRC (Ref: BB/N015762/1) and MRC (MR/L011212/1) grant awarded to ESJR.

## Author contributions

JB – conceptualisation, planning, execution, analysis, interpretation and writing

ER – funding, support for experiment design, interpreting and writing

## Supplementary materials for

**Supplementary Table S1:**
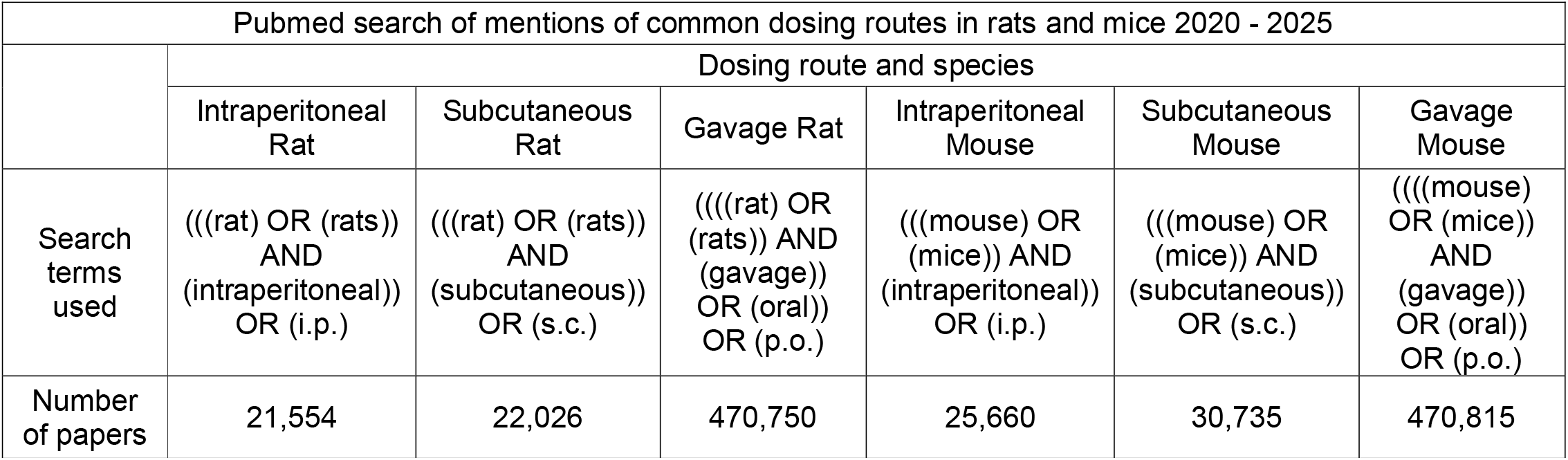
search of common dosing methods in rats and mice.

**Supplementary Table S2:** Condensed Milk Nutritional Information.

| Condensed Milk Nutritional Information |  |
| --- | --- |
| Ingredients | Whole Milk, Sugar |
| Nutrition Information |  |
| Typical Values | Per 100g as sold |
| Energy | 1359kJ / 322kcal |
| Fat | 8.0g |
| of which: saturates | 5.0g |
| Carbohydrates | 55.0g |
| of which: sugars | 55.0g |
| Fibre | 0.0g |
| Protein | 7.5g |
| Salt | 0.26g |

**Figure S1:**
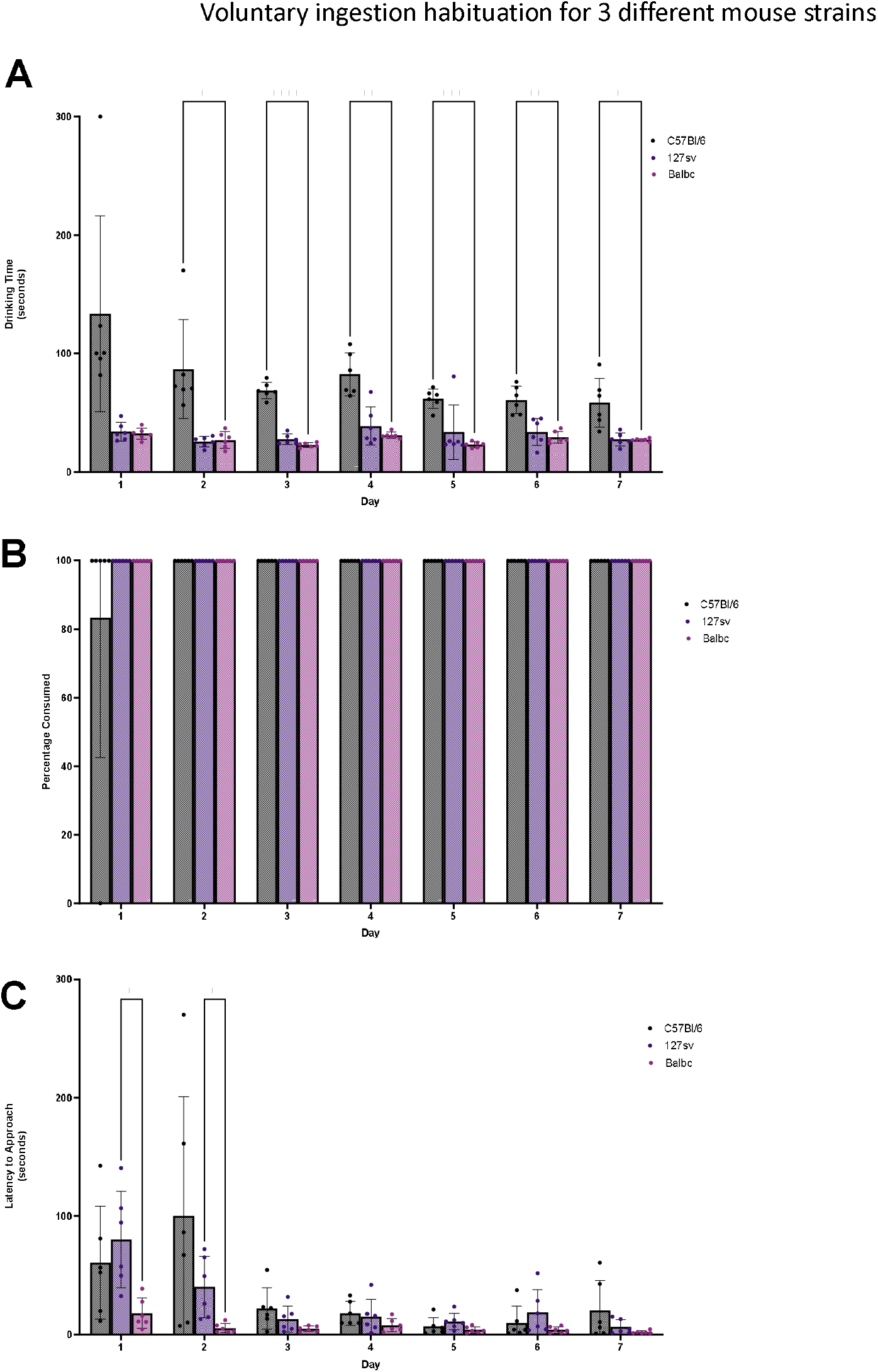
Different strains of mice were presented with 10% condensed milk. **A** C57BL6 mice were significantly slower to drink the entire volume on days 2-7 compared to the 127sv and BalbC mice except on day 5 where they were only significantly slower than the BalbC mice (day 2 p = 0.0445/0.0480, day 3 p = <0.0001/<0.0001, day 4 p = 0.0043/0.0025, day 5 p = 0.0001, day 6 p = 0.0072/0.0018, day 7 p = 0.0373/0.0389. There was also a main effect of time showing a significant decrease in drinking time for all strains over the course of 7 days (p = 0.0404). **B** One C57BL6 mouse did not consume the entire volume from the syringe on day 1. All other mice consumed 100% and from day 2 onwards all mice consumed 100%. **C** 127sv mice were significantly slower to approach the syringe than the BalbC mice on days 1 and 2 (day 1 p = 0.0277, day 2 p = 0.0469). There was also a main effect of time showing a significant decrease in approach time for all strains over the course of 7 days (p = 0.0011). Bars are mean +/-SEM and dots represent individual animals.

### Habituation Protocol for Voluntary Oral dosing Rats and Mice

- A few days before you intend to start the dosing schedule, expose the animals to the vehicle that you will be using in their home cage. An example would be 10% condensed milk and 90% water. They may be reluctant to approach the syringe so, if necessary, the vehicle could be placed on a surface in the cage and left for the animals to investigate in their own time.
- Recommended volumes for mice are ∼0.2ml and rats 0.5ml
- For the following days, present the vehicle in the syringe and use a 5 min cut off time for each animal
- If the animal does not approach the syringe within 5 mins, then leave the vehicle in the cage on a flat surface or in a dish until the next session
- If the animal approaches and drinks from the syringe but does not consume the entire dose within 5 mins then remove syringe as this will encourage quicker drinking next session.
- Once they are happy drinking from the syringe you can start your dosing study. This normally takes about 3-5 days.
- To make up drugs, you can follow your normal formulation protocol but substitute your usual vehicle for the diluted palatable substance. For example, dissolve the drug in water (50% of total vehicle volume) and then add the palatable solution (50% of total vehicle volume) once it has dissolved.
- During the dosing schedule the exact volume required for the animal is drawn up into the syringe.
- If the animals are group housed, then you can either separate the required animal for dosing or dose all animals in the cage at the same time using multiple syringes (rats can become competitive for access to the syringe so usually require separation).

### Palatable solution options

1. Condensed milk – standard dilution 10% but can be increased to 20% for drugs which have a bitter taste to help mask the flavour.
2. Flavoured milkshake - standard dilution 50%. Commercial milkshakes can contain various additives which should be considered in the decision-making process.
3. Fruit juice – standard dilution 50% up to 90%. 4. Saccharin – 1-2%
4. Peanut butter – drug is dissolved in nut oil first and then mixed with peanut butter at a dilution of up to 50%. This can be useful for drugs which don’t dissolve in water.

## References

1. Bonnichsen M D. N.,, Hansen A. The welfare impact of gavaging laboratory rats. Animal Welfare 14, 223–227 (2005). 10.1017/S0962728600029389

2. Balcombe JP, Barnard ND, Sandusky C. Laboratory routines cause animal stress. Contemp Top Lab Anim Sci. 2004 Nov;43(6):42–51. PMID: 15669134.

3. Azar, T., Sharp, J. & Lawson, D. Heart rates of male and female Sprague-Dawley and spontaneously hypertensive rats housed singly or in groups. J Am Assoc Lab Anim Sci 50, 175–184 (2011).

4. Stuart, S. A. & Robinson, E. S. Reducing the stress of drug administration: implications for the 3Rs. Sci Rep 5, 14288 (2015). 10.1038/srep14288

5. Gärtner, K. et al. Stress response of rats to handling and experimental procedures. Lab Anim 14, 267–274 (1980). 10.1258/002367780780937454

6. Gonzales, C. et al. Alternative method of oral administration by peanut butter pellet formulation results in target engagement of BACE1 and attenuation of gavage-induced stress responses in mice. Pharmacol Biochem Behav 126, 28–35 (2014). 10.1016/j.pbb.2014.08.010

7. Walker, M. K. et al. A less stressful alternative to oral gavage for pharmacological and toxicological studies in mice. Toxicol Appl Pharmacol 260, 65–69 (2012). 10.1016/j.taap.2012.01.025

8. Płaźnik, A., Stefański, R. & Kostowski, W. Restraint stress-induced changes in saccharin preference: the effect of antidepressive treatment and diazepam. Pharmacol Biochem Behav 33, 755–759 (1989). 10.1016/0091-3057(89)90466-8

9. LaFollette, M. R. et al. Laboratory Animal Welfare Meets Human Welfare: A Cross-Sectional Study of Professional Quality of Life, Including Compassion Fatigue in Laboratory Animal Personnel. Front Vet Sci 7, 114 (2020). 10.3389/fvets.2020.00114

10. Kaźmierowska, A. M. et al. Rats respond to aversive emotional arousal of human handlers with the activation of the basolateral and central amygdala. Proc Natl Acad Sci U S A 120, e2302655120 (2023). 10.1073/pnas.2302655120

11. Arantes-Rodrigues, R. et al. The effects of repeated oral gavage on the health of male CD-1 mice. Lab Anim (NY) 41, 129–134 (2012). 10.1038/laban0512-129

12. Brown, A. P., Dinger, N. & Levine, B. S. Stress produced by gavage administration in the rat. Contemp Top Lab Anim Sci 39, 17–21 (2000).

13. Germann, P. G. & Ockert, D. Granulomatous inflammation of the oropharyngeal cavity as a possible cause for unexpected high mortality in a Fischer 344 rat carcinogenicity study. Lab Anim Sci 44, 338–343 (1994).

14. Isaksson, I. M., Theodorsson, A., Theodorsson, E. & Strom, J. O. Methods for 17β-oestradiol administration to rats. Scand J Clin Lab Invest 71, 583–592 (2011). 10.3109/00365513.2011.596944

15. Kalliokoski, O., Jacobsen, K. R., Hau, J. & Abelson, K. S. Serum concentrations of buprenorphine after oral and parenteral administration in male mice. Vet J 187, 251–254 (2011). 10.1016/j.tvjl.2009.11.013

16. Küster, T. et al. Voluntary ingestion of antiparasitic drugs emulsified in honey represents an alternative to gavage in mice. J Am Assoc Lab Anim Sci 51, 219–223 (2012).

17. Abelson, K. S., Jacobsen, K. R., Sundbom, R., Kalliokoski, O. & Hau, J. Voluntary ingestion of nut paste for administration of buprenorphine in rats and mice. Lab Anim 46, 349–351 (2012). 10.1258/la.2012.012028

18. Atcha, Z. et al. Alternative method of oral dosing for rats. J Am Assoc Lab Anim Sci 49, 335–343 (2010).

19. Corbett, A., McGowin, A., Sieber, S., Flannery, T. & Sibbitt, B. A method for reliable voluntary oral administration of a fixed dosage (mg/kg) of chronic daily medication to rats. Lab Anim 46, 318–324 (2012). 10.1258/la.2012.012018

20. Diogo, L. N. et al. Voluntary Oral Administration of Losartan in Rats. J Am Assoc Lab Anim Sci 54, 549–556 (2015).

21. Ferguson, S. A. & Boctor, S. Y. Use of food wafers for multiple daily oral treatments in young rats. J Am Assoc Lab Anim Sci 48, 292–295 (2009).

22. The 3Hs Initiative [Internet]. https://www.3hs-initiative.co.uk. x[cited 2024 October 16]. Available from: https://www.3hs-initiative.co.uk/the-3hs

23. Kronenberger, J. P. & Médioni, J. Food neophobia in wild and laboratory mice (Mus musculus domesticus). Behav Processes 11, 53–59 (1985). 10.1016/0376-6357(85)90102-0

24. Modlinska, K. & Stryjek, R. Food Neophobia in Wild Rats (Rattus norvegicus) Inhabiting a Changeable Environment-A Field Study. PLoS One 11, e0156741 (2016). 10.1371/journal.pone.0156741

25. Rodent MDA [Internet]. https://www.rodentmda.ch [cited 2024 October 16]. Available from: https://www.rodentmda.ch

26. Tillmann, S. & Wegener, G. Syringe-feeding as a novel delivery method for accurate individual dosing of probiotics in rats. Benef Microbes 9, 311–315 (2018). 10.3920/bm2017.0127

27. Sjödén, P. O. & Archer, T. Conditioned taste aversion to saccharin induced by 2, 4, 5-trichlorophenoxyacetic acid in albino rats. Physiol Behav 19, 159–161 (1977). 10.1016/0031-9384(77)90174-3

28. Andrews, D. et al. Bitter-blockers as a taste masking strategy: A systematic review towards their utility in pharmaceuticals. Eur J Pharm Biopharm 158, 35–51 (2021). 10.1016/j.ejpb.2020.10.017

29. Coupland, J. N. & Hayes, J. E. Physical approaches to masking bitter taste: lessons from food and pharmaceuticals. Pharm Res 31, 2921–2939 (2014). 10.1007/s11095-014-1480-6

30. Mennella, J. A., Pepino, M. Y. & Beauchamp, G. K. Modification of bitter taste in children. Dev Psychobiol 43, 120–127 (2003). 10.1002/dev.10127

31. Mennella, J. A., Reed, D. R., Roberts, K. M., Mathew, P. S. & Mansfield, C. J. Age-related differences in bitter taste and efficacy of bitter blockers. PLoS One 9, e103107 (2014). 10.1371/journal.pone.0103107

32. De Oliveira Sergio, T., Frasier, R. M. & Hopf, F. W. Animal models of compulsion alcohol drinking: Why we love quinine-resistant intake and what we learned from it. Front Psychiatry 14, 1116901 (2023). 10.3389/fpsyt.2023.1116901

33. De Villiers, M. M. in A practical guide to contemporary pharmacy practice Vol. 3 (ed J.E. Thompson) Ch. 22, 267–276 (Lippincott Williams & Wilkins, 2009).

34. Noorjahan, A., Amrita, B., & Kavita, S. (2014). In vivo evaluation of taste masking for developed chewable and orodispersible tablets in humans and rats. Pharmaceutical development and technology, 19(3), 290–295. 10.3109/10837450.2013.778870

35. Grau-Bové, C., Grau-Bové, X., Terra, X., Garcia-Vallve, S., Rodríguez-Gallego, E., Beltran-Debón, R., Blay, M. T., Ardévol, A., & Pinent, M. (2022). Functional and genomic comparative study of the bitter taste receptor family TAS2R: Insight into the role of human TAS2R5. FASEB journal : official publication of the Federation of American Societies for Experimental Biology, 36(3), e22175. 10.1096/fj.202101128RR

36. Cheney, C. D.,Miller, E.R. Effects of forced flavor exposure on food neophobia. Applied Animal Behaviour Science 53, 213–217 (1997). 10.1016/S0168-1591(96)01160-4

37. Ingram, D. K. Lithium chloride-induced taste aversion in C57BL/6J and DBA/2J mice. J Gen Psychol 106, 233–249 (1982).

38. Prendergast, M. A., Hendricks, S. E., Yells, D. P. & Balogh, S. Conditioned taste aversion induced by fluoxetine. Physiol Behav 60, 311–315 (1996). 10.1016/0031-9384(95)02234-137

39. Hinchcliffe, J. K., Stuart, S. A., Mendl, M., & Robinson, E. S. J. (2017). Further validation of the affective bias test for predicting antidepressant and pro-depressant risk: effects of pharmacological and social manipulations in male and female rats. Psychopharmacology, 234(20), 3105–3116. 10.1007/s00213-017-4687-5

40. Stuart, S. A., Butler, P., Munafò, M. R., Nutt, D. J., & Robinson, E. S. (2015). Distinct Neuropsychological Mechanisms May Explain Delayed-Versus Rapid-Onset Antidepressant Efficacy. Neuropsychopharmacology : official publication of the American College of Neuropsychopharmacology, 40(9), 2165–2174. 10.1038/npp.2015.59

41. A(SP)A. (https://www.gov.uk/government/publications/the-operation-of-the-animals-scientific-procedures-act-1986/the-operation-of-the-animals-scientific-procedures-act-1986-aspa-accessible, 2014).

42. EU Directive 2010/63/EU DIRECTIVE 2010/63/EU OF THE EUROPEAN PARLIAMENT AND OF THE

43. COUNCIL of 22 September 2010 on the protection of animals used for scientific purposes. https://eur-lex.europa.eu/legal-content/EN/TXT/PDF/?uri=CELEX:32010L0063

